# PhageScanner, a flexible machine learning pipeline for automated bacteriophage genomic and metagenomic feature annotation

**DOI:** 10.1101/2023.07.17.549438

**Authors:** Dreycey Albin, Mirela Alistar

## Abstract

Even though bacteriophages are the most plentiful organisms on Earth, many of their genomes and assemblies from metagenomic sources lack protein sequences with identified functions. Most proteins in bacteriophages are structural, known as Phage Virion Proteins (PVPs), but a considerable number remain unclassified. Complicating matters further, conventional lab-based methods for PVP identification are time-consuming and tedious. To expedite the process of identifying PVPs, machine-learning models are increasingly being employed. While existing tools have developed models for predicting PVPs from protein sequences as input, none of these efforts have built software allowing for genomic and metagenomic as input. In addition, there isn’t a framework available for easily curating data and creating new types of models. In response, we introduce PhageScanner, an open-source platform that streamlines data collection, model training and testing, and includes a prediction pipeline for annotating genomic and metagenomic data. PhageScanner also features a graphical user interface (GUI) for visualizing annotations on genomic and metagenomic data. We also introduce a BLAST-based classifier that outperforms ML-based models (achieving an F1 score of 94% for multiclass PVP detection and 97% for binary PVP detection) and an efficient Long Short-Term Memory (LSTM) classifier. We showcase the capabilities of PhageScanner by predicting PVPs in six previously uncharacterized bacteriophage genomes. In addition, showing the utility of the framework, we create a new model that predicts phage-encoded toxins within bacteriophage genomes.

## Introduction

Bacteriophages (or phages), recognized as the most prolific organisms on Earth [1, 2], play an integral role in shaping bacterial ecology [3, 4]. They have also emerged as a potential therapeutic alternative against infections caused by antibiotic-resistant bacteria [5]. An essential precursor to the widespread use of phage therapy depends on developing a comprehensive understanding of phage structural components, their function, and other proteins that may be toxic to humans [6]. This importance is compounded by the fact many bacteriophage proteins within assembled genomes and metagenomic data cannot be assigned a function due to the lack of other similar proteins [7]. These structural proteins, often referred to as bacteriophage virion proteins (PVPs), play a role in processes such as bacterial recognition, bacteriophage proliferation, and the overall bacteriophage life cycle - which necessitates thorough exploration. Identifying these proteins traditionally involves high-throughput experimental strategies, such as mass spectrometry [8] and protein arrays [9–11]. However, these techniques can be labor-intensive and time-consuming [12, 13].

Consequently, there is a growing interest in leveraging computational techniques to expedite and simplify the identification of these proteins [12, 13].

The computational identification of PVPs first gained traction in 2011 when a group utilized feed-forward neural networks for binary classification of proteins as either structural or non-structural [14]. The selection of networks was accuracy-driven, and this measure was further improved by implementing an ensemble of 160 networks that were used together to classify proteins through a majority voting scheme [14]. To validate this computational approach, the researchers employed Transmission Electron Microscopy (TEM) to examine proteins identified using this method, including those forming capsids and tail fibers.

Since that seminal study, numerous computational approaches have been developed, each contributing updated classification methods with improved accuracy. Until 2019, these innovations primarily targeted the binary classification of phage proteins as either PVPs or not, utilizing traditional machine learning (ML) algorithms like random forests (RFs) [15], Support Vector Machines (SVMs) [16], Naive Bayes (NB) [17], or ensemble methods combining these classifiers. A shift in focus occurred in 2020 when Cantu et al. released “PhANNs”, a software that applied multiclass classification to identify the specific type of PVP among 10 different classes [18]. This change represented a significant advancement as it shifted from simple PVP identification to the determination of precise types of PVP. Two years later, DeepPVP, proposed by Fang et al., built upon this multiclass approach by employing a convolutional neural network (CNN) for PVP type identification [19]. Leveraging the PhANNs dataset, this work enhanced prediction performance for multiclass PVP prediction and demonstrated promising results for binary PVP prediction as well.

While existing approaches have substantially advanced the identification of bacteriophage PVPs, there are still potential improvements to be made in relation to the datasets used, the frameworks for model testing and training, and the practical implementation of predictors in bioinformatics applications. Many methodologies have utilized Uniprot to source ground truth proteins [15–17], whereas recent tools have relied exclusively on proteins obtained from Entrez [18, 19]. Furthermore, tools not using pre-established test sets have had to re-perform database querying and the creation of testing and training splits. These variances in model creation and testing could result in inconsistencies between studies. Additionally, these studies have not extended their predictors’ application to genomic or metagenomic data, DeePVP being a notable exception though not as a part of the software [19].

Our research extends upon prior studies by introducing PhageScanner, an open-source tool that empowers users to predict PVPs from various inputs such as reads, genomes (or assembled contigs), and direct proteins. The tool’s comprehensive machine learning and prediction pipeline is depicted in Figure 1. PhageScanner automates data curation and model training processes, and the software also houses a prediction pipeline compatible with trained models. This tool addresses past research shortcomings by providing a platform that facilitates the creation and testing of protein feature prediction models (binary or multiclass) using proteins sourced from both UniProt and Entrez. We also incorporate a BLAST classifier as a predictor, allowing for a direct comparison of previous ML-based classifiers to a bioinformatic solution using sequence alignment. Moreover, PhageScanner integrates a workflow for annotating Open Reading Frames (ORFs) within genomic and metagenomic data, along with a graphical user interface (GUI) for scraping proteins of interest from the predictions.

**Fig 1.**
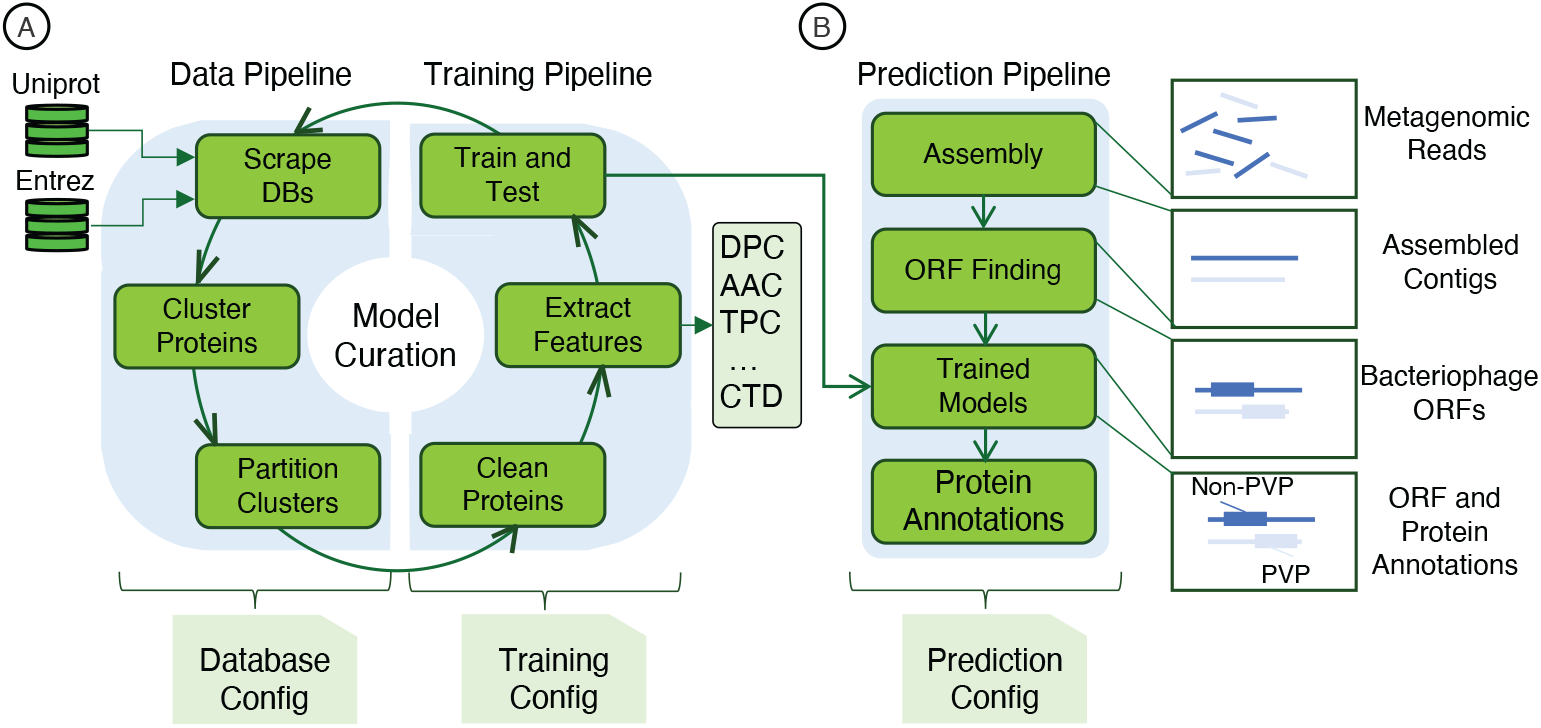
Overview of the PhageScanner Machine Learning (ML) Pipeline. The ML pipeline is composed of three distinct yet interconnected sub-pipelines: (1) The Data Pipeline, (2) The Training Pipeline, and (3) The Prediction Pipeline. Each pipeline is designed to be executed conveniently from the command line, and features a customizable configuration file. (A) The process of model curation is split into two stages. The “Data Pipeline” is responsible for curating data through adapters connected to a protein database. Subsequently, the “Training Pipeline” undertakes the extraction of features and initiates model training as defined by the corresponding configuration file. (B) The “Prediction Pipeline” employs a trained model, generated by the Training Pipeline, to identify Open Reading Frames (ORFs) within metagenomes, genomes, or proteins directly.

Lastly, we show the utility and flexibility of the pipeline by creating models to predict bacteriophage-encoded toxins, which may help in ensuring the safety of future phage therapies. The source code for the pipelines and the GUI, as well as pre-trained models, are available at https://github.com/Dreycey/PhageScanner.

## Design and Implementation

### Reconfigurable framework

The PhageScanner framework is comprised of three underlying pipelines: the (1) Data Pipeline, (2) Training Pipeline, and (3) Prediction Pipeline (Fig 1). Each pipeline has a corresponding configuration file that can be modified for customized usage. The Data Pipeline retrieves protein data from Entrez and/or Uniprot [20, 21], clusters this data using sequence similarity, and separates protein clusters into partitions for downstream training. The Training Pipeline extracts vectorized features from the database-retrieved protein sequences to train downstream models (Fig 1A). Lastly, the Prediction Pipeline uses parameterized models from the Training Pipeline to predict protein classes from reads, genomes or proteins directly (Fig 1B). Together PhageScanner provides a novel framework – the first two pipelines enable the automation of creating bacteriophage protein annotation models, and the final pipeline allows end-users to apply these models to annotate proteins within genomic or metagenomic datasets.

## Data pipeline

PhageScanner was designed to integrate multiple protein databases and curate proteins for training downstream machine learning models. This process is performed within the Data Pipeline, which begins with gathering proteins from Uniprot [20] and Entrez [21]. Here each protein annotation (i.e. prediction class) is assigned a query to Uniprot and/or Entez to gather proteins corresponding to that annotation, each resulting in a fasta file. While previous work used either Entrez or Uniprot, this approach is novel in that it utilizes both databases to retrieve proteins. After retrieval, proteins are then clustered using CD-HIT [22] using an identity threshold defined in the corresponding configuration file. Following this organization, each protein cluster, along with its individual members, is divided into *K* partitions for performing *k*-fold cross-validation [23]. Here, *K* signifies the total partitions made for training and testing; specifically, one partition is allocated for testing, while the remaining K-1 partitions are dedicated to training.

When applying this methodology to collect proteins corresponding to the 10 PVP classes as outlined in earlier studies [18, 19], we discern several key observations.

Initially, at a clustering threshold of 90%, most clusters are comprised of fewer than 400 proteins (Fig 2A). Furthermore, when the identity threshold for clustering is more strict (i.e. decreased), we observe a declining trend in the number of total clusters (Fig 2B). This trend is most notable when the identity drops from 100% to 90%, resulting in the greatest reduction in the number of clusters. These findings suggest a high prevalence of proteins belonging to smaller clusters and highlights the existence of many unique proteins within the queries. In addition, there is a substantial difference in the initial count of proteins between Uniprot and Entrez. Specifically, Entrez holds a significantly higher count of proteins per class compared to Uniprot (Fig 2C). Overall, using a combination of Uniprot and Entrez along with clustering offers an advantage in creating larger datasets for downstream model training.

**Fig 2.**
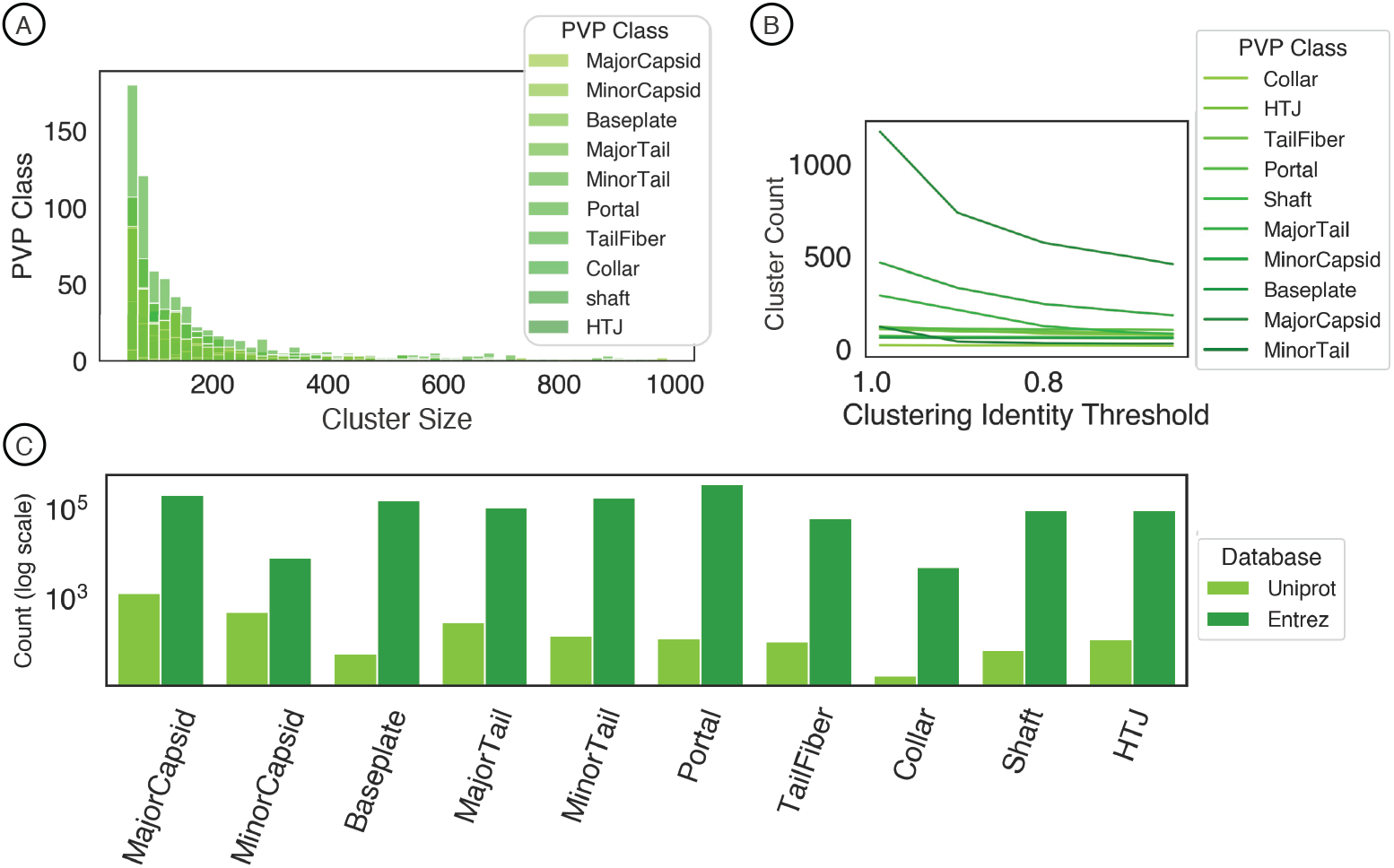
Database retrieval and clustering. (A) When the proteins are clustered at an identity threshold of 90% using CD-HIT, all classes of the proteins show a higher frequency of small cluster sizes, while few have clusters larger than 400 proteins. (B) The number of clusters at different identity thresholds using CD-HIT. (C) The count of proteins from both Uniprot and Entrez before clustering.

### Training pipeline

The training pipeline utilizes the partitioned proteins from the Database Pipeline to extract features from the proteins and subsequently train and test models. The process begins with the cleaning of proteins by removing all non-canonical amino acids from the sequences [24]. Thereafter the features specified in the configuration file of the training pipeline are extracted from each protein. These extracted features then serve as inputs for training downstream models using the k-fold cross-validation partitions created in the Data Pipeline [13]. The separation of the Training Pipeline from the Database Pipeline offers flexibility, enabling end-users to iteratively test models without needing to redownload or partition the proteins.

### Feature extraction

The feature extraction segment of the pipeline allows end-users to experiment with various feature extraction techniques commonly applied for PVP identification and other protein-focused models [13]. In PhageScanner, we employ a factory design pattern [25], a software design approach that facilitates the simple combination of different extracted features into a more comprehensive feature vector. In essence, each feature method extracts a specific feature from the provided protein, and the resulting feature vectors are subsequently concatenated to form a more extensive vector (Equation 1; see S1 Text for feature descriptions). The advantage of this approach is twofold. Firstly, it enables developers to effortlessly incorporate new feature extraction methods. Secondly, it offers users the flexibility to determine which feature to utilize for each model within the configuration file.

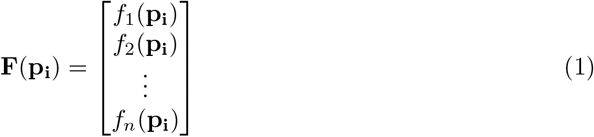

We evaluated each feature extraction method for accuracy and timing using a baseline one-vs-all logistic regression classifier. Initial findings demonstrated that tripeptide and dipeptide frequency features were most accurate for multiclass PVP prediction, with TailFiber proteins proving the hardest to predict (Fig 3A). When we combined features into a single concatenated vector, as shown in Eq. 1, the combination of di- and tri-peptide frequencies was the most accurate combinatorial-predictor (Fig 3B). Other feature combinations also yielded baseline models with high F1 scores, highlighting PhageScanner’s capacity for feature investigation. Notably, di- and tri-peptide feature extraction required the most time, due to the large feature vector size (8400 elements) - a crucial consideration for extensive metagenomic dataset analysis (see S1 Text for table).

**Fig 3.**
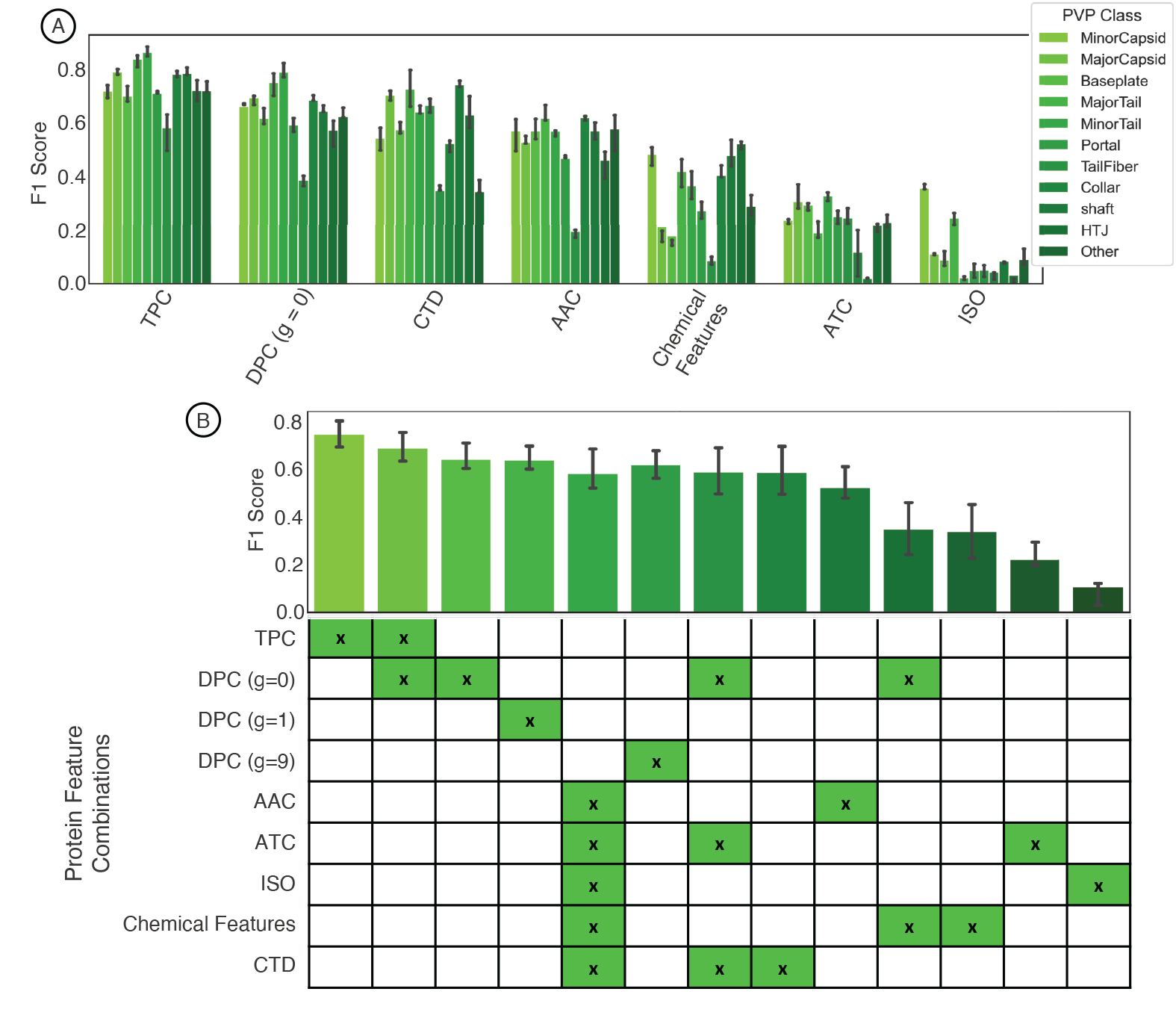
Protein feature extraction testing. (A) The F1 score for different standalone features using a baseline logistic regression (one-vs-all) classifier. Colors correspond to the specified phage virion protein class. (B) Mean F1 scores among all classes for different feature combination approaches.

### Model testing and selection

Once features are selected, the Training Pipeline evaluates models, each with its own set of pre-defined feature vectors, specified in the configuration file. PhageScanner has a variety of models defined in the software but allows for end users to easily add new models. We test the two types of models frequently used for PVP prediction: (1) multiclass prediction (for 10 PVP proteins as defined in [18]), and (2) binary prediction (determining if a protein is PVP or not, as per [14]). For each case, PhageScanner includes a BLAST classifier that takes the top-scoring protein alignments from BLAST and selects the highest-scoring class as the label, akin to a K-nearest neighbor classifier. Additionally, using Keras [26], we added a long-short-term memory (LSTM) deep learning model to PhageScanner as these are designed for sequential information. These methods are then compared to reimplementations of other classification models described in the literature [13]. However, we do not use the software provided by other tools as PhageScanner can directly incorporate models, and the other software tools do not use data from both Uniprot and Entrez.

In the context of multiclass PVP prediction, we compared the performance of the BLAST classifier and LSTM against a reimplementation of PhANNs’ feed-forward neural network (FFNN) and DeePVP’s convolutional neural network (CNN). As a baseline, we also employed a logistic regression one-vs-all classifier. Upon evaluation, it was clear that the BLAST classifier surpassed other PVP prediction methods in accuracy (94%), leaving the FFNN (86%) and LSTM (82%) as the next highest performing models (Fig 4A-top). However, while the BLAST classifier demonstrated the best accuracy, its inference time was significantly higher than the other prediction models, as expected (Fig 4A-bottom). For the binary prediction model, which determines if a protein is a PVP in general, the BLAST classifier outperformed the learning methods with a mean f1 score of 94% (Fig 4B-top). Both PVP-SVM and LSTM showed comparable performance, with both yielding average F1 scores of 91%. All models surpassed the baseline classifier that uses logistic regression. As seen with the multiclass prediction models, the BLAST classifier took the longest for inference, while all other classifiers ran in less than a second on the test datasets (Fig 4B-bottom).

**Fig 4.**
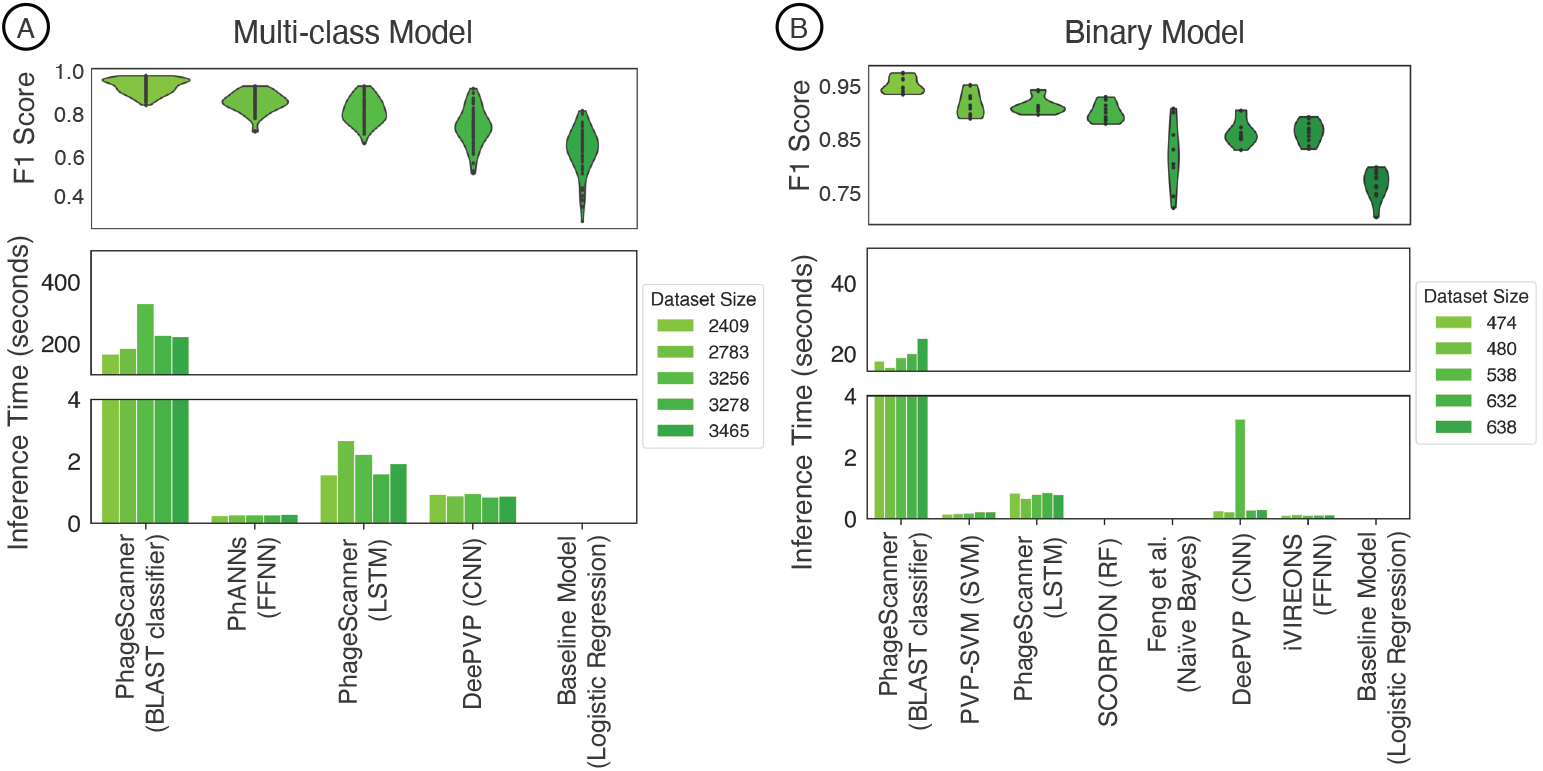
Testing performance among models for different classification tasks. (A) In the context of multiclass PVP prediction, the upper panel displays the F1 score across various models, while the lower panel illustrates the inference time of different multiclass classifiers. Inference timing was performed on dataset sizes, post clustering, of 2409, 2783, 3256, and 3465 proteins. (B) For binary PVP prediction, the upper panel demonstrates the F1 score across different models, whereas the lower panel showcases the inference among various binary classifiers. Inference timing was performed on dataset sizes, post clustering, of 474, 480, 536, 632, and 838 proteins.

**Fig 5.**
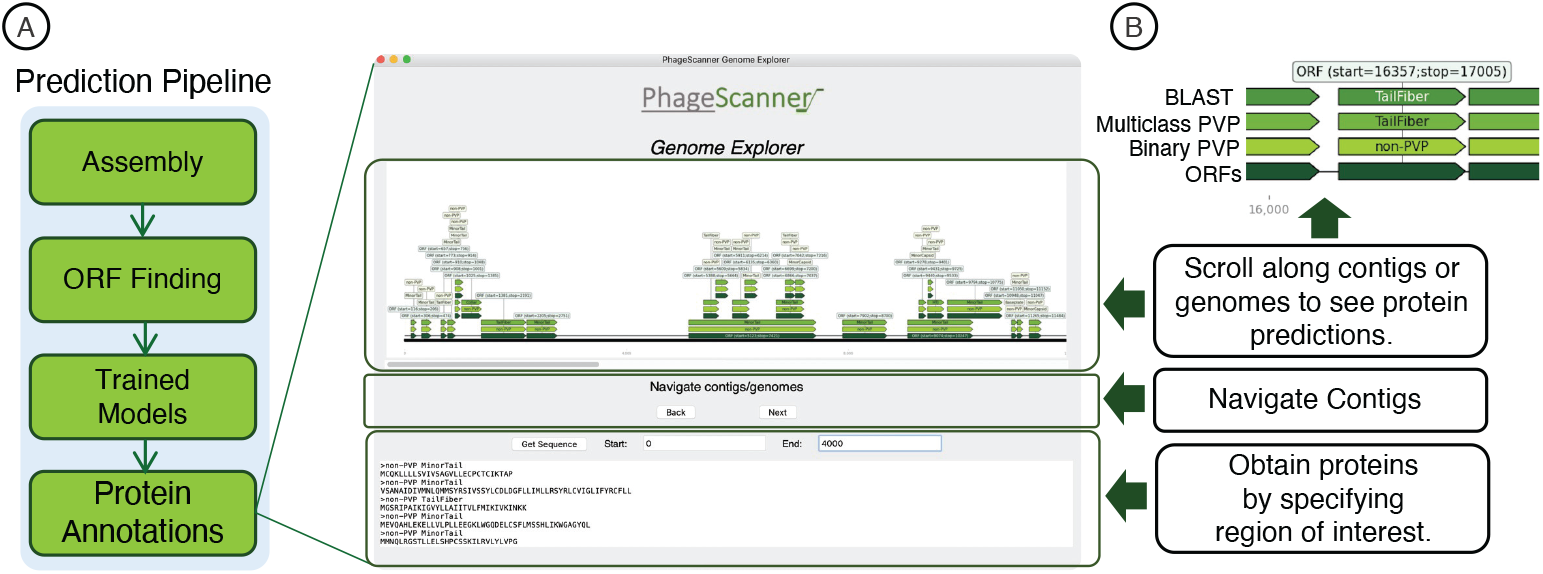
Graphical User Interface (GUI) for visual assessment of genomic and metagenomic annotations. (A) The prediction pipeline produces a comma-separated (CSV) summary of features found for the different genomic and metagenomic annotations. This output can be explored using a provided, open-source, graphical user interface (GUI). (B) The GUI provides a method for end users to easily navigate across different annotations while cross-referencing different annotation models with BLAST. The contigs and/or genomes can be scrolled across, and the lower panel of the GUI allows for retrieving proteins specified within a genomic range for further analysis.

### Prediction pipeline and GUI

Once a model has been trained and tested, it can be deployed in the Prediction Pipeline to annotate open reading frames (ORFs) originating from genomic and metagenomic data. If metagenomic sequencing data is used as input, the pipeline starts by assembling this data using the Megahit assembly software [27]. The output of this process consists of longer assembled continuous segments, termed contigs. Of note, the assembly step is skipped if genomes are used as input. Due to its bacteriophage-focused algorithm, PHANOTATE is then used to identify ORFs from the contigs or genomes [28]. These ORFs are converted into their corresponding protein products, which are subsequently transformed into predefined feature vectors by each model specified in the configuration file for the Prediction Pipeline. These feature vectors then serve as input for the pre-trained models to annotate the proteins. Many models used within the pipeline, allowing for cross-referencing predictions and allowing for different types of protein feature annotations at once. For example, models can be used to predict if an ORF consists of a PVP protein and/or a toxin. This presents the first work applying these PVP prediction models as a foundation for annotating bacteriophage genomes (as opposed to only using proteins as input).

The Prediction Pipeline generates a comma-delimited output file that can be analyzed using the provided graphical user interface (GUI). Constructed using the TKinter library from Python’s standard library [29], the GUI starts by using the DNA Features Viewer library [30] to create feature-specific visualizations of each ORF along the length of a contig or genome. Each model used in the Prediction Pipeline will be given a track along the contig/genome. Once these input images are generated, the GUI is then populated with the output CSV file and the corresponding image directory, enabling a visual exploration of annotated contigs and genomes. An additional feature of the GUI is its lower frame, which enables listing all proteins within a specific section of the genome or contigs when a particular region is selected. This interactive setup allows users to swiftly extract proteins of interest from the contigs or genomes processed by the prediction pipeline.

## Results and discussion

As outlined, PhageScanner incorporates a BLAST classifier and an LSTM model for predicting PVP classes. While the LSTM model performs well overall, it struggles to accurately classify certain PVP classes, as illustrated by the performance gap between it and the BLAST classifier (91% vs 94%). A deeper exploration into the LSTM’s performance per PVP class revealed a few challenging categorizations (Fig 6A). The model frequently misclassifies Portal proteins as Minor Capsid proteins, and confuses Tail Fiber proteins with Collar proteins. Given their similar functions, such misclassifications are expected and also seen in the work of PhANNs [18]. Furthermore, a significant number of proteins are falsely identified as the negative class “Other”. This issue may be rooted in PVPs being contained within the protein dataset corresponding to the “Other” class and can potentially be addressed by refining the database queries. As per our analysis, we additionally see the LSTM struggles most with the “Other” category and “Tail Fibers” class when comparing aggregated F1 scores across each PVP class (Fig 6B).

**Fig 6.**
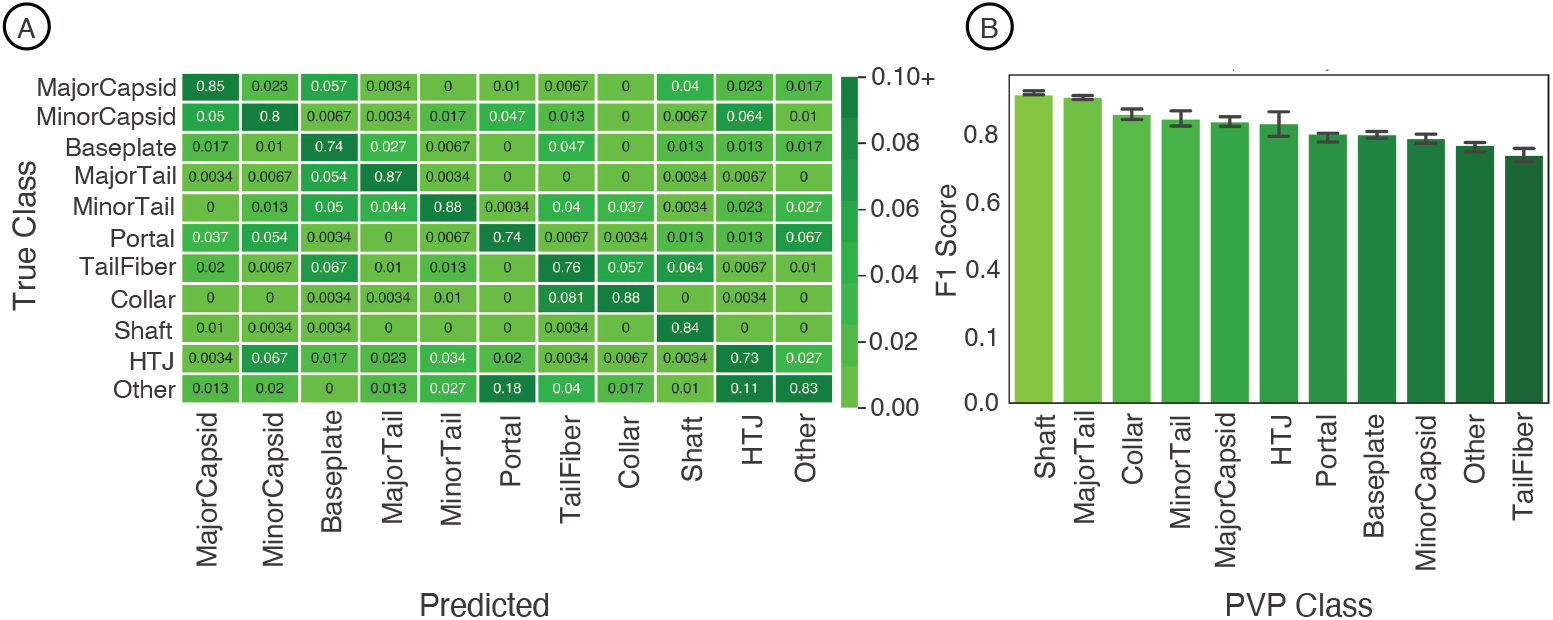
Analysis of Multiclass Performance of PhageScanner on Phage Virion Proteins. (A) A confusion matrix displaying the frequency from the ground truth class (rows) to the predicted class (columns). As most off-diagonal entries fall below 90%, the cell color spectrum is adjusted to range between 0 and 0.1. (B) The F1 score per class for the PhageScanner LSTM model, with the highest scoring classes arranged in descending order of F1 score.

To showcase the utility of PhageScanner, we used the trained LSTM model and the BLAST classifier to predict open reading frame (ORF) annotations on uncharacterized bacteriophage genomes. We used six genomes sourced from the open-access archive of 25,152 bacteriophage genomes assembled in April 2023 as part of the Infrastructure for a Phage Reference database [31]. The genome accessions we analyzed were AC171169, BK010471, GU339467, MF417929, MH552500, and MH616963. Each genome was input into the prediction pipeline as a multi-fasta file, with subsequent analyses performed on the output prediction CSV file.

A fascinating observation from our analysis was that most of the PVP classes tended to be located in specific regions across each genome, with the Major Capsid class typically appearing at the start of the genomes (Fig 7A). Interestingly, when we examined the occurrence of different PVP classes within each genome, we did not find a consistent pattern (Fig 7B). We note this lack of consistency might be due to our limited sample size of six randomly selected genomes. A tally of each class per genome revealed that while Major Capsid did not occur frequently, other PVPs like TailFibers were commonly observed throughout the genomes (see S1 Text for PVP counts per genome). Most importantly, these results illustrate the ability of PhageScanner to be used as a tool for annotating PVPs within bacteriophage genomes.

**Fig 7.**
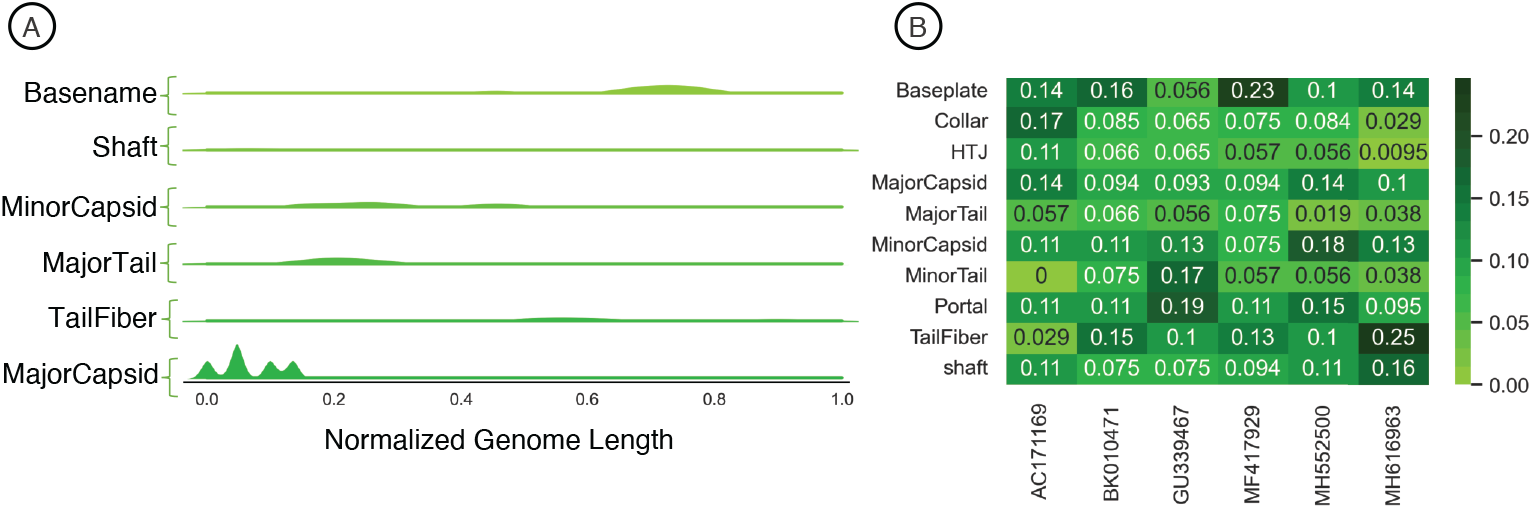
Analysis of Phage Virion Proteins (PVPs) in Uncharacterized Genomes. (A) A Kernel Density Estimation (KDE) distribution for each predicted class, aggregated along the normalized length of all genomes. (B) The frequency of different PVP classes within each genome, with the frequency calculated on a per-genome basis.

## 0.1 Finding Bacteriophage Toxins

While bacteriophages can be used as an effective alternative to antibiotics for treatment, there are potential safety concerns that need to be assessed. There are many known instances where bacteriophages aid in the pathogenicity of certain bacteria, assisting in virulence that leads to human disease. Examples of this phenomenon occurring in nature include *Streptococcus pyogenes* and *Vibrio cholerae* developing toxigenicity after the induction of toxin-encoding bacteriophages [6, 32]. Here we use PhageScanner to develop models that can predict the presence of toxic proteins from bacteriophage genomes.

Verheust et al. curated a list of bacteriophage-encoded toxins associated with human diseases. We used this refined set to train machine-learning models to predict toxins within bacteriophage genomes. Specifically, nine phage-encoded toxins were defined in this set, and we established Uniprot queries for downloading each type. We also defined a negative class (i.e. non-toxin), consisting of all reviewed bacteriophage proteins that are not associated with toxicity according to the gene ontology database [33] (see S1 Text for all queries). Following this process, we collated a set of 857 toxin proteins for the positive class (i.e. Toxin), and 1989 non-toxin proteins for the negative class (i.e. non-Toxin).

While developing a prediction model for toxins, we tested the BLAST classifier, a support vector machine (SVM), a long short-term memory (LSTM) network, and a feed-forward neural network (FFNN). We also used logistic regression as a baseline model for comparison. All models displayed lower F1 scores in comparison to the models predicting PVPs (80% compared to 94%) (Fig 8A). Of note, false positives were observed more frequently than false negatives, which is desirable as false negatives would cause potentially toxic proteins to be missed (Fig 8B).

**Fig 8.**
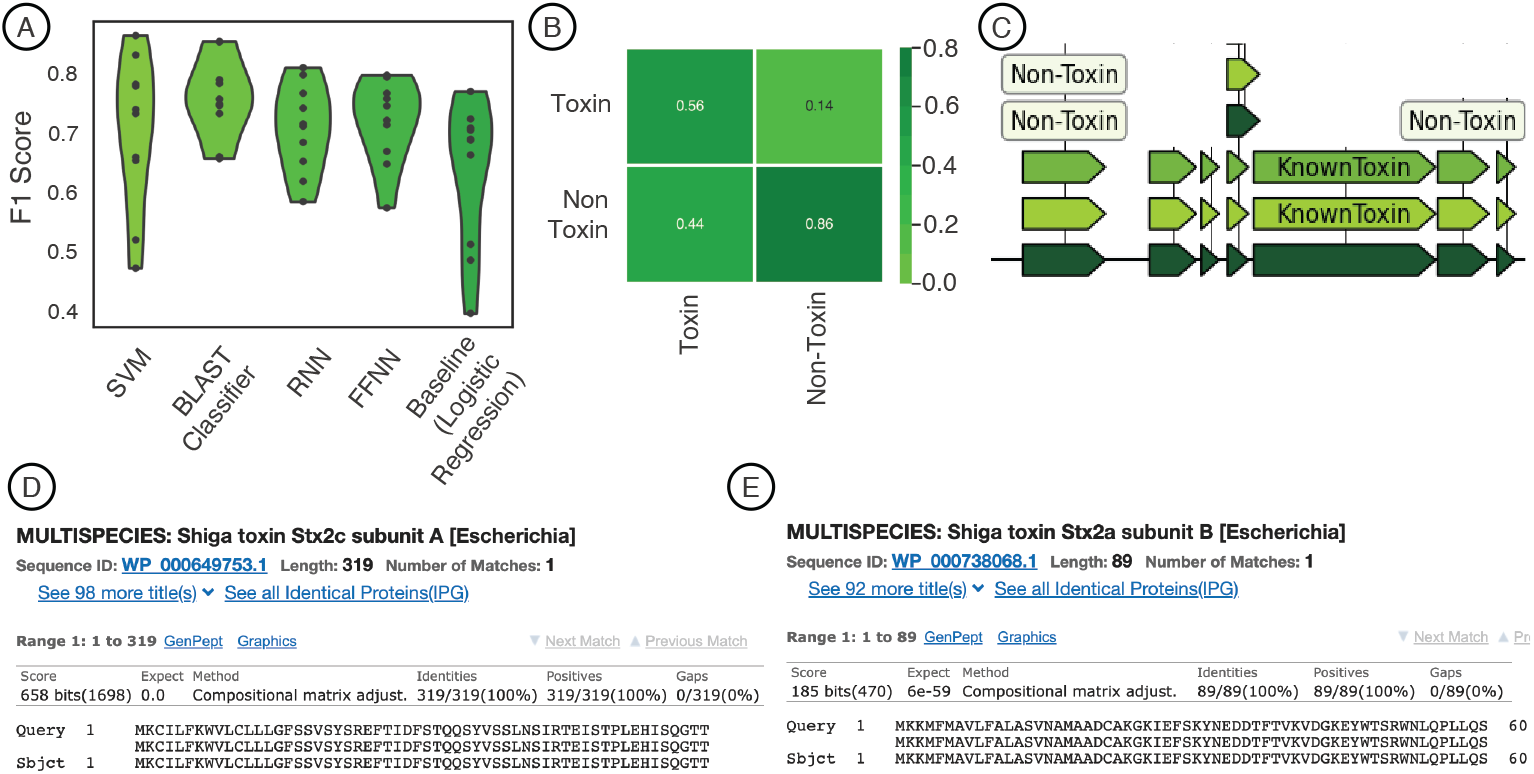
Using PhageScanner to create models for predicting phage-encoded toxins within bacteriophage genomes. (A) F1 Scores for several models in predicting if a give protein is a phage-encoded toxin. The models tested include a support vector machine (SVM), the BLAST classifier, a long short-term memory (LSTM) network, and a feed-forward neural network (FFNN). (B) Confusion matrix for the BLAST classifier with true labels on in the rows and predicted labels in the columns. (C) Screenshot from the GUI showing Stx1 (Shiga toxin 1) as classified being a toxin by both the FFNN and BLAST classifier. (D) BLAST web server screenshot validating the Stx1 (Shiga toxin 1) translated protein. (E) BLAST web server screenshot validating the Stx2 (Shiga toxin 2) translated protein.

To test the prediction models, we use the FFNN and BLAST classifier in the Prediction Pipeline on *Escherichia* phage 933W (NCBI ID: NC 000924.1). coding Shiga toxins 1 and 2 (Stx1 and Stx2), this phage contributes to the pathogenic potential of *Escherichia coli* O157:H7 [34, 35], thereby establishing itself as a robust positive control for assessing the models and approach. Feeding the final prediction CSV into the GUI allowed for visually scanning the genome for toxin-focused feature annotations (Fig 8C), showing the Stx1 and Stx2 ORFs as predicted known toxins by both the FFNN and BLAST classifier. To validate the predictions, the proteins translated in the expected region of the toxins were validated using the BLAST web server [36] for both Stx1 and Stx2 (Fig 8D-E).

## Availability and Future Directions

PhageScanner is an open-source project that you can freely access and use, as it is licensed under the GNU General Public License. If you’re interested, the project is hosted on GitHub and can be found at this link: https://github.com/Dreycey/PhageScanner. Notably, the GitHub page also offers options for subscribing or unsubscribing to the PhageScanner mailing list.

Going forward, our plan is to focus on enhancing the performance of PhageScanner with the goal of making it highly efficient for use on high-performance computing systems. Since PhageScanner is an open-source software, we also anticipate that community feedback will play a crucial role in steering its future development.

## Supporting information

**S1 Text. Supplemental material and tables**. The supplemental file includes software usage, tables showing performance metrics for different feature combinations (S1 Table), counts of phage virion proteins (PVPs) found in each uncharacterized bacteriophage genome (S2 Table), and information about queries for phage-encoded toxins (S3 Table).

## Acknowledgments

This work utilized the Summit supercomputer, which is supported by the National Science Foundation (awards ACI-1532235 and ACI-1532236), the University of Colorado Boulder, and Colorado State University. The Summit supercomputer is a joint effort of the University of Colorado Boulder and Colorado State University.

## Notes

### Competing Interest Statement

The authors have declared no competing interest.

